# hdnom: Building Nomograms for Penalized Cox Models with High-Dimensional Survival Data

**DOI:** 10.1101/065524

**Authors:** Nan Xiao, Qing-Song Xu, Miao-Zhu Li

## Abstract

**Summary:** We developed hdnom, an R package for survival modeling with high-dimensional data. The package is the first free and open-source software package that streamlines the workflow of penalized Cox model building, validation, calibration, comparison, and nomogram visualization, with nine types of penalized Cox regression methods fully supported. A web application and an online prediction tool maker are offered to enhance interactivity and flexibility in high-dimensional survival analysis.

**Availability:** The hdnom R package is available from CRAN: https://cran.r-project.org/package=hdnom under GPL. The hdnom web application can be accessed at http://hdnom.io. The web application maker is available from http://hdnom.org/appmaker. The hdnom project website: http://hdnom.org.

## 1 Introduction

The dimensionality of collected data in clinical studies for complex diseases, such as cancer, is growing exceedingly fast. It is analytically challenging for researchers to elucidate the relationship between the most influential factors and patient survival outcomes. In practice, penalized Cox regression models (Gui and Li, 2005) are usually favored among other machine learning methods for excellent interpretability. Such penalized linear models achieve simultaneous variable selection and effect size estimation, while still have considerable predictive performance. For example, researchers have built such models for prostate cancer prognosis prediction (Halabi *et al.*, 2014) based on high-dimensional clinical measurements. On the other hand, high-throughput genomic and epigenomic characterization data have also been analyzed with such methods, to identify key molecular-level signatures contributing to patient survival probability in colon cancer (Zhang *et al.*, 2013). Notably, a recent DREAM challenge also proposed the same problem on the quantitative prediction of survival outcomes with high-dimensional clinical trial data (Abdallah *et al.*, 2015). Therefore, there is an emerging need for building such survival models with high-dimensional data. However, creating a validated survival model ready for clinical use is not a trivial process. It requires intensive computational efforts and extensive validation procedures. Unfortunately, many of the available software packages for survival modeling usually focus on model fitting, lacking necessary features to support model evaluation and visualization, thus created a significant gap between theory and practice.

To fill such gap for high-dimensional survival modeling, we started the hdnom project, which offers a freely available R package, a web application, and an online prediction tool maker. To the best of our knowledge, hdnom is the first free and open-source software package which streamlined the workflow of high-dimensional Cox model building, validation, calibration, comparison, and nomogram visualization. With hdnom, clinical researchers can easily build prognostic models, evaluate model performance, and prepare publication-quality figures in a reproducible way.

## 2 Methods

### 2.1 Available Penalized Cox Regression Methods

For high-dimensional linear models, a substantial number of penalized estimation methods have been proposed during the past decade, and many of them have been successfully applied in high-dimensional Cox regressions. Based on their maturity and popularity, nine types of penalized estimation methods were selected to be supported in hdnom, including lasso, adaptive lasso, fused lasso, elastic-net, adaptive elastic-net, SCAD, Snet, MCP, and Mnet. The selected methods have different theoretical properties and variable selection effects (Fu and Xu, 2012; Xiao and Xu, 2015). They are considered to be representative enough to accomplish specific variable selection tasks for most types of clinical and genomic data. It could be difficult to have a comprehensive understanding of the methods or choose an appropriate one sometimes, see supplement 1 for an introduction to all the supported methods, and brief guidelines on how to choose from these methods.

Most of the penalized estimation problems supported by hdnom are solved by efficient optimization procedures, such as coordinate descent algorithms. For penalized Cox models, all the listed methods have one or more parameters to tune. To minimize the burden of tedious manual inspection, most of the parameter tuning steps are automated using internal cross-validation in hdnom when fitting the model. Usually, users are only expected to specify their preferred parameter grids for a minimal number of parameters. If not specified, the accuracy of the models built with the default settings is often still reasonable. In hdnom, the parameter tuning procedures can run in parallel efficiently, which further reduced the time required when building survival models for large-scale datasets.

### 2.2 Model Validation, Calibration, and Comparison

Rigorous model evaluation is critical for predictive modeling (Simon *et al.*, 2011). Three resampling-based methods, namely bootstrap, cross-validation, and repeated cross-validation are supported in hdnom for model validation and model calibration. For model validation, three types of time-dependent AUC estimators are available as the evaluation metrics (supplement 1). Empirical quantiles of the metrics generated from the resampling are also given. Such summary statistics could be informative as confidence measures for estimation accuracy. For model calibration, Kaplan-Meier survival curves and log-rank test are supported for the risk groups derived by calibration. With hdnom, the built survival models can be validated and calibrated using internal data (by resampling) and using (independent) external datasets. To help the users gain more insight into the empirical model performance when choosing from the available penalized estimation methods, functions for model comparison by validation or calibration are provided. Eventually, publication-quality figures can be generated for all model validation, calibration, and comparison results, to facilitate further visual analytics.

### 2.3 Nomograms for Penalized Cox Model Visualization

Although it is not essential to make predictions using graphs directly today, nomogram is still a simple yet effective graphical representation of linear models. Nomograms have been extensively used by clinicians when interpreting Cox regression models in clinical research and publications. All the baseline hazard functions were estimated using the Breslow estimator in hdnom. The *rms* package (Harrell, 2015) was utilized to create the nomogram visualizations for all types of models, and we used ordinary least squares to make the approximations needed for creating the mapping between the model coefficients and the coordinates in the nomogram.

### 2.4 Web Application and Application Maker

For users with less programming experience, we also offer an interactive web application (hdnom.io) which supports all features available in the R package. A reproducible report in PDF/HTML/Word formats can be generated and downloaded after necessary modeling steps are finished in the application, with the essential information for reproducing the modeling process. With the hdnom web application, users can also make direct survival probability predictions for new samples after a model was built. Furthermore, for users who prefer to publish their own model-based online prediction applications, such as (Rudloff *et al.*, 2010), a web application maker is provided. After downloading the built model object file from the hdnom web application, the application maker will help the users produce and serve their unique online prediction applications efficiently.

## 3 Conclusions

We anticipate that the survival models, visualizations, and disease-specific online prediction tools created by hdnom can help clinicians assess individualized risk factors, predict the likely outcomes of treatment, and make data-driven decisions on follow-up patient-tailored strategies. See supplement 2 for a hands-on tutorial for the R package, and supplement 3 for a reference manual with examples for all built-in functions in hdnom.

## Acknowledgements

We thank Prof. Matthew Stephens from Department of Human Genetics and Department of Statistics at the University of Chicago, Prof. Samuel Volchenboum, Dr. Jorge Andrade from Center for Research Informatics at the University of Chicago, Dr. Svetlana Ukraintseva from Duke University, and Dr. Liang Cheng from Indiana University for their suggestions on improving the software quality. We thank our collaborators at Peking University Cancer Hospital, Peking University People's Hospital, and Sun Yat-sen University Cancer Center for testing the web application and offering useful feedbacks. We also thank Dr. Yihui Xie and Dr. Garrett Grolemund from RStudio Inc. for providing help on hosting the web application.

